# Resurrecting the ancient glow

**DOI:** 10.1101/778688

**Authors:** Yuichi Oba, Kaori Konishi, Daichi Yano, Hideyuki Shibata, Dai-ichiro Kato, Tsuyoshi Shirai

**Affiliations:** Department of Environmental Biology, Chubu University 487-8501, Japan; Graduate School of Bioagricultural Sciences, Nagoya University 464-8601, Japan; Graduate School of Science and Engineering, Kagoshima University 890-0065, Japan; Department of Bioscience, Nagahama Institute of Bio-Science and Technology, 526-0829, Japan

## Abstract

The colour of firefly bioluminescence is primarily determined by the structure of the enzyme luciferase^1^. To date, firefly luciferase genes have been isolated from over 30 extant species producing light ranging in colour from deep-green to orange-yellow. We have reconstructed ancestral firefly luciferase genes and characterised the enzymatic properties of the recombinant proteins in order to predict ancestral firefly light emission. Results showed that the synthetic luciferase for the last common firefly ancestor exhibited green light. All known firefly species are bioluminescent in the larval stages^2^, with a common shared ancestor arising approximately 100 Mya^3^. Combined, our findings propose within the Cretaceous forest the common ancestor of contemporary fireflies emitted green light, most likely for aposematic display from nocturnal predation.

---

Over the centuries the bioluminescence of fireflies has attracted much attention as a charming seasonal sight, particularly in Asia, and recently as a useful diagnostic tool in the biomedical sciences^4^. It is proposed that firefly bioluminescence originated as an aposematic warning display toward predators, and later acquired a role in sexual communication for many firefly species^2^.

Fireflies belong to the beetle family Lampyridae; composed of 7 subfamilies containing around 2,000 recognized species around the world^2^. Luminescent beetles are additionally found in the families Phengodidae, Rhagophthalmidae and Elateridae. It has been considered that Phengodidae and Rhagophthalmidae are sister groups of the Lampyridae (‘cantharoid’ group) and share a common origin of bioluminescence; conversely bioluminescence in Elateridae evolved independently of the cantharoid group^5^.

The molecular system of bioluminescence is shared among these four families: the chemical structure of the luminescent substrate, D-luciferin, is considered identical for all luminescent beetles, and the luminescence reaction is catalyzed by homologous luciferases (> 48% AA identity) in the presence of O_2_, ATP, and Mg^2+^ (Fig. 1a)^5^. Despite the commonality in enzymatic reaction and components, luminescence colour can vary widely between species. The European glowworm *Lampyris noctiluca*, for example, emits green light, the North American Big Dipper firefly *Photinus pyralis* yellow-green light, and the Japanese lesser firefly *Luciola parvula* orange-yellow light. The differences in luminescence colour is considered to be the consequence of evolutional strategies for warning predators and attracting mating partners more effectively^2, 6^.

**Figure 1.**
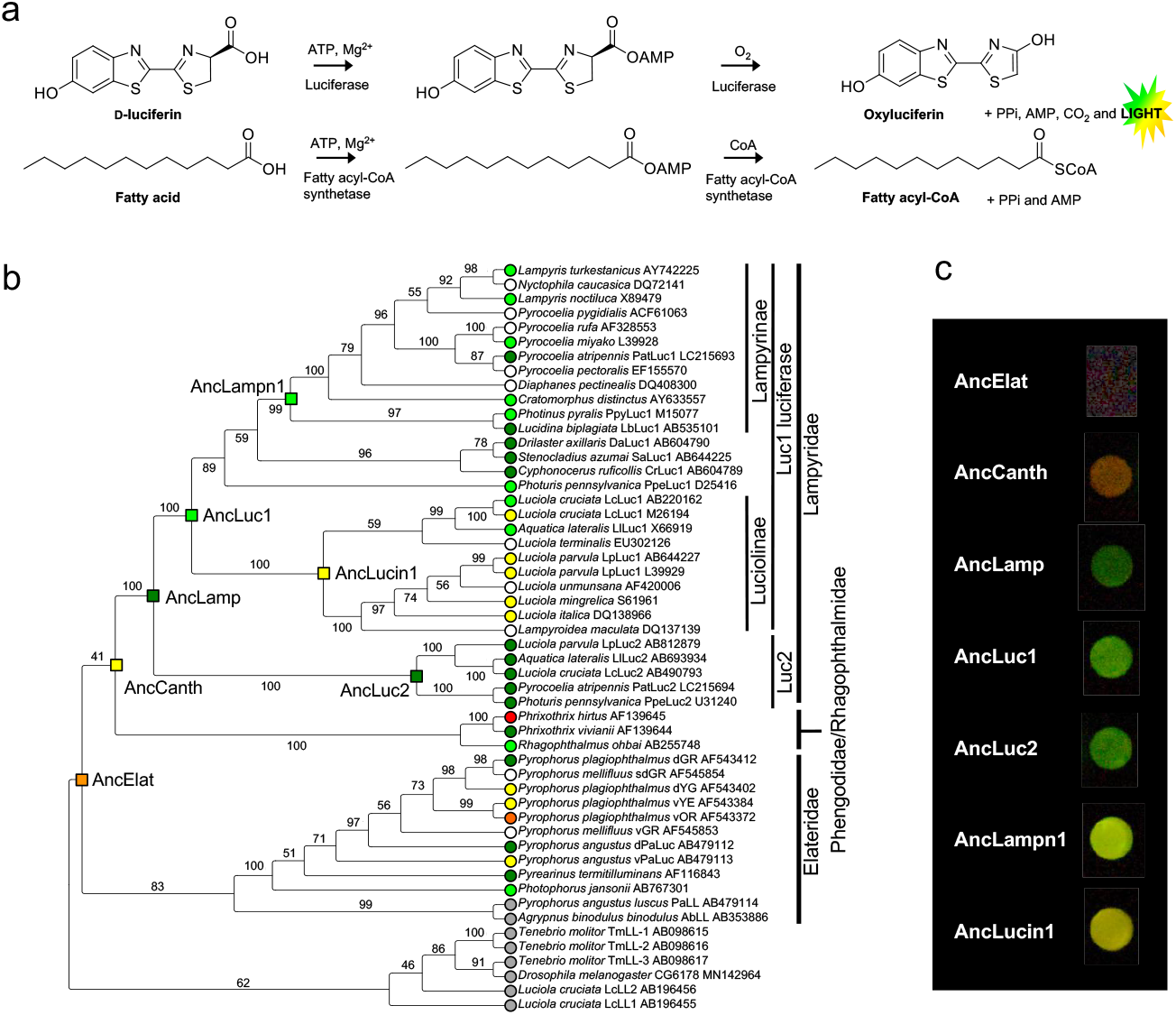
**a**, Coleopteran bioluminescence reaction. **b,** Molecular phylogeny of luciferases and related enzymes. The leaf nodes are labelled with species name, protein name and GenBank accession number. Branches are labelled with bootstrap probability (1,000 reconstructions). The resurrected ancestral nodes are shown as a square. The leaf and resurrected ancestral nodes are indicated with *in vitro* luminescent colours (green, yellow-green, yellow, orange, or red) judged by the luminescence maximum value (Table 1, also text). Luciferases without spectral data and fatty acyl-CoA synthetase (non-luciferase) are denoted with white and grey circles, respectively. **c,** Photographs of the luminescence of seven ancestral luciferases in 96-well plate. The camera exposure time for each well is uneven to avoid the changes in coloration by overexposure.

**Table 1.** Reconstructed ancestral luciferases.

| Ancestral luciferase | Position | Median divergence age (Mya) [95% HPD] | Av. P.P.<sup>a</sup> | $\lambda_{\text{max}}$ (nm)<sup>b</sup> and colour |
| --- | --- | --- | --- | --- |
| AncElat | Elateroidea ancestor | 115.85 [84.82 - 145.26] | 0.915 | 594 (orange) |
| AncCanth | ‘Cantharoids’ ancestor | 102.55 [71.06 - 132.83] | 0.915 | 584 (yellow) |
| AncLamp | Lampyridae ancestor | n.a. | 0.935 | 548 (green) |
| AncLuc1 | Luc1 ancestor | 69.59 [44.57 - 96.19] | 0.938 | 550 (yellow-green) |
| AncLuc2 | Luc2 ancestor | 69.59 [44.57 - 96.19] | 0.949 | 548 (green) |
| AncLampn1 | Lampyrinae Luc1 ancestor | 26.23 [22.91 - 66.83] | 0.984 | 553 (yellow-green) |
| AncLucin1 | Luciolinae Luc1 ancestor | 12.18 [0.50 - 35.09] | 0.968 | 563 (yellow) |

<sup>a</sup>Av. P.P., the average posterior probability of the amino acid residues
<sup>b</sup> $\lambda_{\text{max}}$ (nm), luminescence spectral maximum of the purified recombinant protein at pH8.0

Fossil records of fireflies are limited, but a single adult male firefly was mined from Burmese amber dating about 100 Mya, which, surprisingly, exhibited an obvious photophore structure on the abdominal segment^7^. This strongly suggested that the ancestral firefly of the Cretaceous period was already bioluminescent and used the light emission for some purpose. Naturally the colour of ancestral bioluminescence is not evident from fossils records, making it difficult to predict the original function of the firefly light.

The colours of light in fireflies are regulated by the luciferase structure and neither by the luciferin molecule, nor by the effects of colour filters (like the blue-transmission filter of the hatchetfish photophore), nor by fluorescent substances (like *Aequorea* GFP in jellyfish). For this reason, *in vivo* firefly luminescence colours match principally those of the *in vitro* luminescent reaction of the luciferase with D-luciferin^8, 9^, thus the evolutionary history of firefly bioluminescence is traceable by using the sequence data of extant luciferases.

Based on these premises, we recreated putative ancestral firefly luciferase gene sequences using the maximum likelihood method of ancestral reconstruction (Fig. 1b)^10^, and experimentally characterized the enzymatic properties including the luminescence colour. The luminescence spectra of fireflies are basically unimodal with wide half bandwidth ca. 60 nm, thus the light colour can be approximated by the *λ*max value. Therefore, in this study we report the colouration of the light produced by luciferin-luciferase reaction as a visual description ‘green’ for *λ*max 520-549 nm, ‘yellow-green’ for *λ*max 550-559 nm, ‘yellow’ for *λ*max 560-584 nm, and ‘orange’ for *λ*max 585-619 nm, which are almost in agreement with our visual perception.

Tree topologies of the molecular phylogeny of luciferase genes were consistent with nuclear/mitochondrial gene trees^2^. To trace back the evolution of firefly luciferase properties, we targeted seven key ancestral genes corresponding to the ancestral luciferase of Elateroidea (AncElat), the ancestral luciferase of ‘cantharoids’ (AncCanth), the ancestral luciferase of Lampyridae (AncLamp), the ancestral Luc1-type luciferase of Lampyridae (AncLuc1; genomic analyses had shown that fireflies possess two different types of luciferase gene, Luc1-type and Luc2-type, generated by gene duplication occurring at the basal position of the lineage in Lampyridae)^5, 8, 11^, the ancestral Luc2-type luciferase of Lampyridae (AncLuc2), the ancestral Luc1-type luciferase in Lampyrinae, a major subfamily of Lampyridae (AncLampn1), and the ancestral Luc1-type luciferase of Luciolinae, another major subfamily (AncLucin1) (Fig. 1b; Table 1; Extended Data Fig. 1). Monophyly of all ancestral nodes are supported with >99% bootstrap values, except for AncElat (62%). The average posterior probability of the amino acid residues in the seven reconstructed ancestral luciferases were sufficiently high, by ranging from 0.984 to 0.915 (Table 1; Extended Data Fig. 2). Although 2 to 2l sites were estimated rather uncertainly, most of such sites distributed distant from the active sites^1^ of luciferase with a few exceptions (Extended Data Fig. 3).

**Figure 2.**
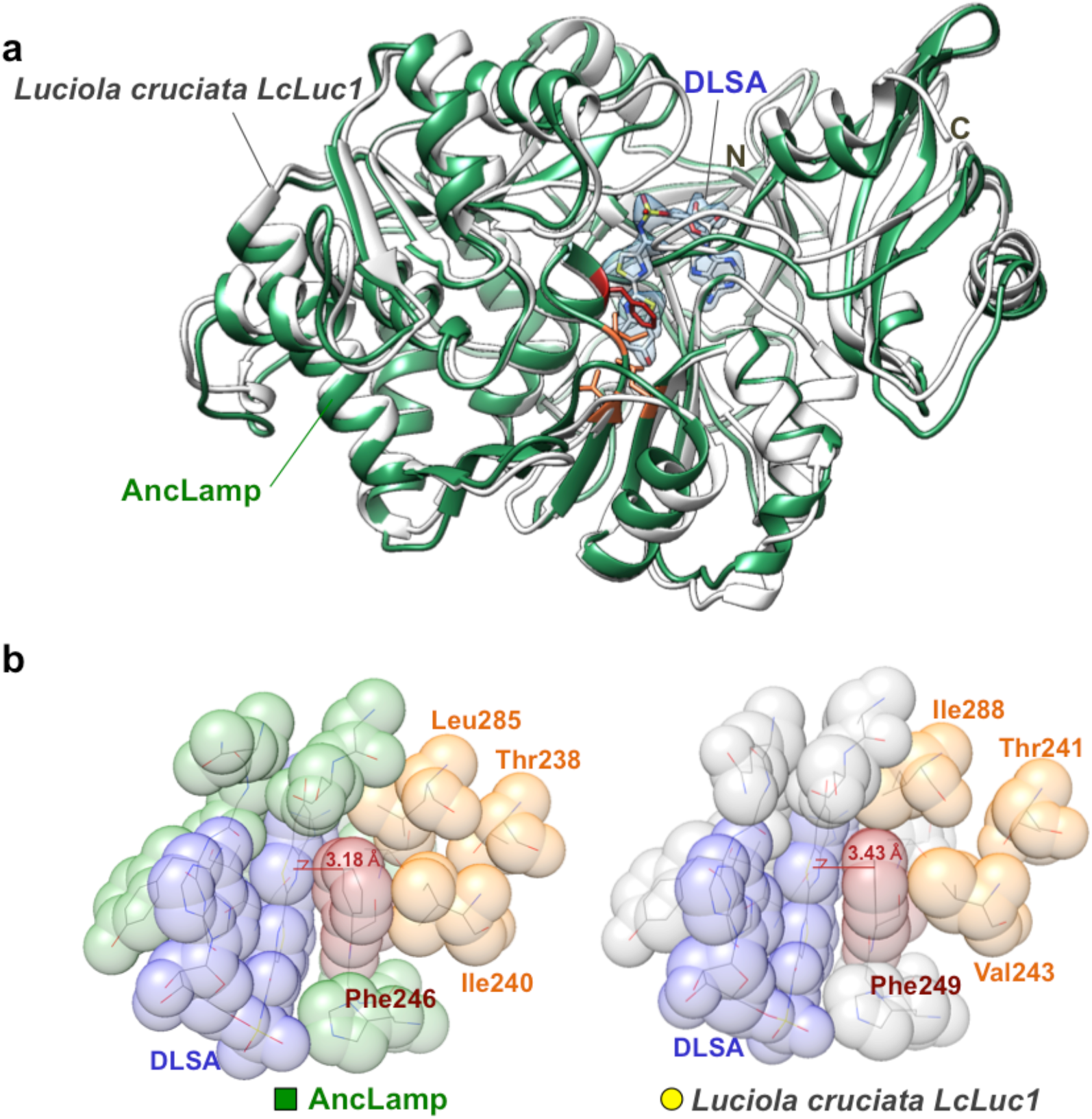
Structure of AncLamp. **a,** The structure of AncLamp – DLSA complex (green) is superposed on that of *Luciola cruciata* LcLuc1 – DLSA complex (PDB 2d1s, grey). DLSA is represented as stick model. The all ligand omitting *Fo* – *Fc* map (contoured at 6 *σ*) is shown in blue. The positions of N- and C-terminals are indicated. **b,** Close up view of the substrate binding site of AncLamp (left) and LcLuc1 (right) in van der Waals model. The different residues between two proteins, i.e. Ile240 (Val243 in LcLuc1) and Leu285 (Ile288), and those in close contact Phe246 (Phe249) and Thr238 (Thr241) are shown labelled. The distances from C*ε*1 atoms of Phe246 (249) to DLSA are indicated in red.

**Figure 3.**
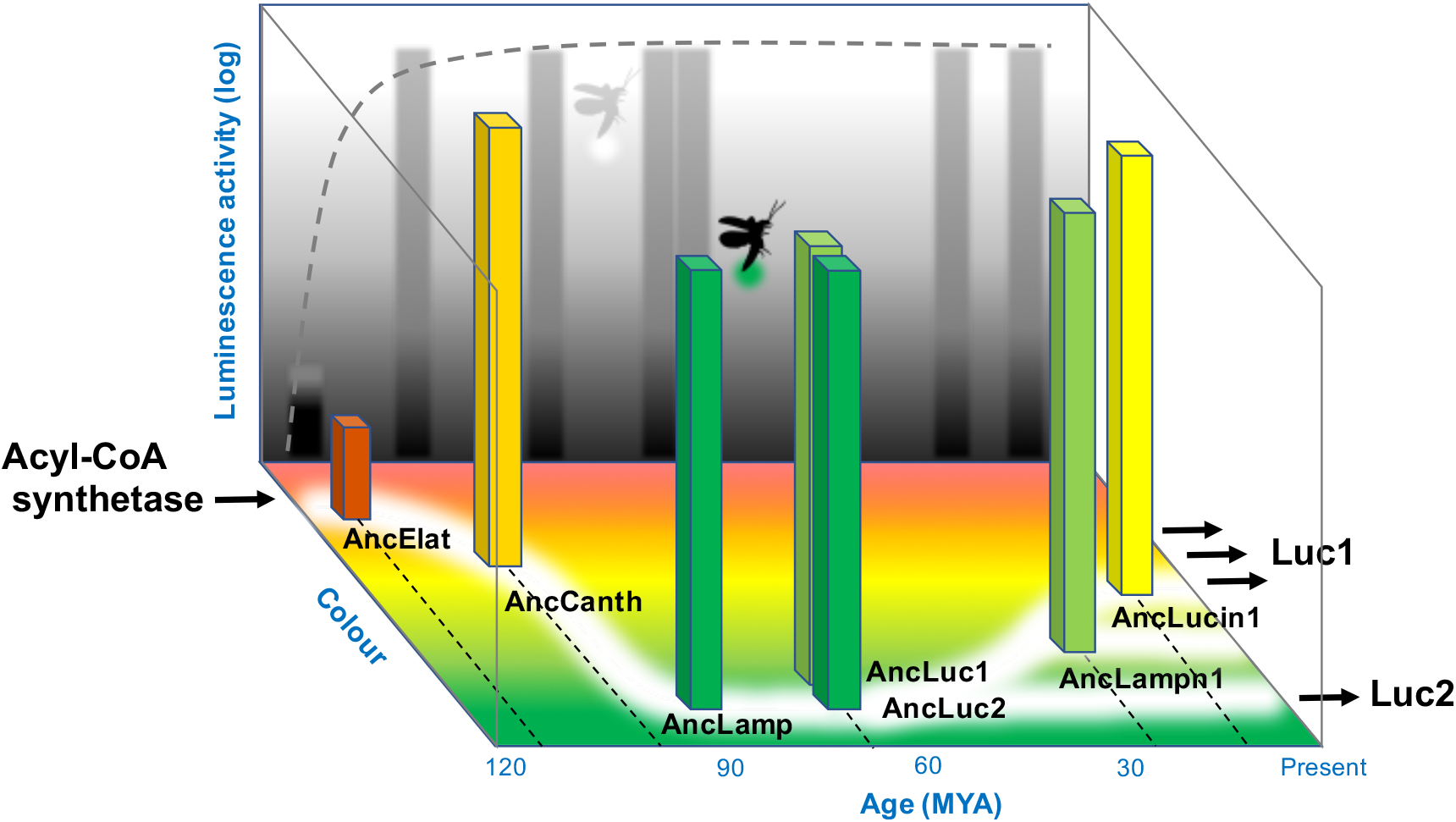
Scheme of the evolution of firefly’s bioluminescence with age (horizontal axis), luminescence activity (vertical axis) and colour z-axis) of ancestral luciferases.

The seven codon-optimized ancestral gene sequences were synthesized and cloned into an expression vector. The recombinant proteins expressed in *E. coli* were purified using a cobalt chelating column (Extended Data Fig. 4). The luminescence assay with D-luciferin showed that six out of seven ancestral luciferases exhibited luminescence activities comparable to extant luciferases (*Luciola cruciata* LcLuc1, LcLuc2 and *P. pyralis* PpyLuc1), but the activity of AncElat was at a trace level (Extended Data Fig. 5). The lack of luminescence activity in AncElat can be rationalized as follows. Position 346 of AncElat (a key residue for luciferin binding^1^) was predicted to be a non-polar Leu (with posterior probability of 0.674). This residue has side chains that cannot interact with the luciferin molecule *via* hydrogen bonding, conversely in all extant luciferases and other ancestral luciferases a polar Ser or Cys residue is found at position 346. Furthermore, this position is Leu in luciferase homologues CG6178 in *Drosophila* and AbLL in non-luminous click beetle, wherein the luminescence activities are negligible^12, 13^. We previously demonstrated that a single mutation of Leu at this position to Ser recovered the significant luminescence activity in both CG6178 and AbLL^13^.

Measurement of the luminescence spectra for seven ancestors showed that: AncElat and AncCanth emit in the orange-red region (*λ*max 594 nm and 584 nm respectively); AncLuc1, AncLuc2 and AncLamp emit in the green region (*λ*max ≤ 550 nm), which is most blue-shifted in seven ancestors; AncLampn1 and AncLucin1 showed a yellow-green to yellow (*λ*max 553 nm and 563 nm), respectively (Table 1, Fig. 1c, Extended Data Fig. 6). These results suggested that: the last common ancestor of ‘cantharoid’ lineage emitted reddish light; the last common ancestor of Lampyridae emitted green luminescence; and after gene duplication, red-shifting occurred in Luc1-type luciferase from green to yellow-green, yellow and orange-yellow during the evolution of the lineages in Lampyrinae and Luciolinae; while Luc2-type luciferase took over the ancestral green light in the extant species^9, 11^.

Firefly luciferases are bifunctional enzymes; in addition to luciferase activity they possess an acyl-CoA synthetic (ACS) activity to various fatty acids^14, 15^; consequently it has been hypothesized firefly luciferase originated from a fatty acyl-CoA synthetase^5^. We measured ACS activities of the seven ancestral luciferases using the substrate lauric acid. Results showed ACS activity of AncElat is 10 times greater than other ancestral and extant luciferases (Extended Data Fig. 7).

These results are consistent with the present evolutionary scenario of luminous beetles. A substantial lack of luminescence activity in AncElat explains the previous conclusion based upon genome analysis^5^, suggesting a parallel origin of bioluminescence in ‘cantharoid’ and Elateridae lineages. High ACS activity of AncElat support the evolutionary ‘trade-off’ model of enzyme neofunctionalization^13, 16^. Reddish luminescence of AncElat and AncCanth fits the hypothesis that ‘protoluciferase’ in beetles, which have a less developed active site environment, emitted in red light^17^. Lastly, green luminescence in AncLamp fit the current hypotheses of firefly biochemists and ecologists, which assumes the luminescence colour of the common ancestor to be green^6, 18–20^ (see more discussion in Supplementary Information).

In order to elucidate the molecular basis of blue-shifted luminescence in AncLamp, the crystal structure was determined in complex with the reaction-intermediate analog 5’-*O*-[*N*-(dehydroluciferyl)-sulfamoyl]adenosine (DLSA) or D-luciferin/ATP by X-ray crystallography to a 2.1 Å or 1.7 Å resolution, respectively (Fig. 2a, Extended Data Fig. 8). AncLamp showed high levels of structural conservation to LcLuc1 (PDB 2d1s), one of the closest extant luciferases showing 79% sequence identity to AncLamp. The previous structural study of LcLuc1 revealed that the steric constraint on luciferyl-moiety was one of the most significant factors for luminescence wavelength; the Ser286Asn mutant of this protein caused a red-shift in wavelength, and the crystal structure indicted that the mutation induced a conformation change in a substrate-binding residue, Ile288, making a space for thermal relaxation of luciferyl moiety^1^. The active sites of AncLamp are generally conserved with LcLuc1 but positions 240 and 285 have Ile and Leu substitutions, respectively (Extended Data Fig. 1). These substitutions appeared to affect the conserved Phe246 conformation eliminating the luciferyl moiety space. The distance from the C*ε*1 atom of Phe residue to the thiazole plane of DLSA is 3.43 Å and 3.18 Å in the LcLuc1 and AncLamp structures, respectively (Fig. 2b). Thus, the blue-shifted luminescence in ancestral luciferase can also be explained by the spatial constraint on the luciferyl moiety (see more discussion in Supplementary Information).

Molecular clock analyses suggested that the origin of the Lampyridae was dated to the mid-Cretaceous^3^. This is consistent with the estimation of geological age of AncCanth of the present study, which suggested 102.55 Mya with 95% HPD of 71.06 to 132.83 Mya (Table 1 and Extended Data Fig. 9).

Green light emission has been observed in the larval stages of the extant diurnal fireflies and also the egg and pupal stages of nocturnal fireflies which, as adults, unlike diurnal fireflies^21^, emit light from their lanterns for sexual communication^9, 11, 22^. Green luminescence is actually common in a number of terrestrial luminous organisms, such as luminous mushrooms and millipedes^23^. The function of the luminescence in fungi is still unclear^24^; as fireflies and millipedes are known to possess distasteful toxins, they are considered to use luminescence as a warning signal^25, 26^. Due to potential nocturnal predators that were maximally sensitive to green light, such emissions producing aposematic displays may have be advantageous for fireflies at night^19, 20^.

In conclusion, we predict that the most ancestral fireflies emitted green light for aposematic display purposes, and some species evolved to use the light as a secondary role for sexual communication, shifting to a more yellowish light (Fig. 3). We believe that our results demonstrated the practical usefulness of the ancestral gene reconstruction methods^10^ successfully reenacting visual scenarios of the lost world.

## Acknowledgements

The authors thank Drs. Victor B. Meyer-Rochow (Andong University, Korea and Oulu University, Finland) and John C. Day (Centre for Ecology & Hydrology, NERC, UK) for critical comments and discussions, and Dr. Shusei Kanie (Nagoya University, Japan) to provide the synthetic DLSA. We are grateful to the Japan Synchrotron Radiation Research Institute (JASRI) for support at the synchrotron facilities at SPring-8. This work was partly supported by Grants-in-aid for scientific research from the Ministry of Education, Science and Culture of Japan (JP17H01818) and Basis for Supporting Innovative Drug Discovery and Life Science Research (BINDS) from the AMED, Japan (19am0101111j0003) to T.S., and JST CREST (JPMJCR16N1) to Y.O.

## Contributions

Y.O., K.D. and T.S. conceived and designed the study. K.K., D.Y., H.S., K.D. and T.S. carried out the molecular and biochemical experiments. T.S. performed the ancestral sequence analyses and crystallography. Y.O., K.D., and T.S. wrote the manuscript and edited the manuscript, based upon discussions with all authors.

## Supplementary Information

### Materials and Methods

#### Ancestral luciferase sequence prediction

The initial set of amino acid sequences was retrieved from the GenBank, Refseq, and UniProt databases. The sequences were aligned by using MAFFT v7.309, and manually refined with the XCED v8a.3.93 program^1^. The final alignment contained a total of 53 sequences (Extended Data Table 2). The topology of phylogenetic tree was inferred based on the amino acid sequences of the extant proteins with the neighbor-joining (NJ) method on the JTT matrix^2, 3^. Then, the tree topology was manually refined so that it was coherent to the currently accepted species phylogeny by referring to the literatures^4, 5^.

The phylogenetic tree and the alignment were applied to PAML v4.5 to infer the ancestral sequences^6^. The empirical model with the JTT matrix was used for the substitution model. No site partition was defined, and substitution rate was uniform over the sites^7, 8^. The ancestral sequences were verified by reconstructing molecular phylogenies to determine if the inferred sequences were connected to the corresponding nodes with zero evolutionary distance. The average posterior probabilities over the amino acid sites for each ancestor are shown in Table 1. The most probable amino acid sequences are aligned in Extended Data Fig. 1. The distributions of site posterior probabilities for each ancestral luciferase are shown in Extended Data Fig. 2. The numbers and distribution of less certainly identified residues, for which the posterior probability is less than 0.1 higher from that of the second most probable amino acid, are shown in Extended Data Fig. 3.

#### Ancestral luciferase geological age estimation

The geological ages of the ancestral luciferases were estimated with the Bayesian Markov chain Monte Carlo (MCMC) method using BEAST v1.8.4. A total of 63 18S rRNA gene sequences of Elateriformia species were aligned by using MAFFT v7.309, and the alignment was manually corrected. A total of 665 aligned sites were selected by avoiding the sites on or near the undetermined or gapped sites. The alignment and the phylogeny based on that of luciferases were submitted for a MCMC calculation. The general time reversible (GTR) substitution model with gamma categories of 4 was adapted. The fossil date 152 Mya (s.d. 10 Mya) of TMRCA (the most recent common ancestor) of Elateroidea^4^ was used for the reference, and the dates of the nodes were estimated on the uncorrelated relaxed clock model. A total of 10^8^ MCMC iterations were performed and the 95% highest posterior density (95% HPD) interval was obtained excluding the first 10^7^ iterations as a burn-in process. The trajectories were analyzed with TRACER v1.6.0 and FIGTREE v1.4.2. The results of ancestral node dating using the Bayesian MCMC method demonstrated that the date of the reference node (Elateroidea common ancestor) has adequately converged and distributed normally around the expected date with mean of 150.7 Mya and 95% HPD from 131.15 to 170.46 Mya based on 87588 effective sample size (ESS). The estimated geological ages of ancestral luciferases are shown in Table 1.

#### Expression and purification of recombinant proteins

Full ORF of the ancestral sequences were synthesized (Fasmac, Kanagawa, Japan) with *E. coli* optimized codon use, and ligated in frame into the expression vector pCold-ZZ-P-X^9^. The vectors were subcloned into *E coli* BL21 (DE3) pLysS (Promega, Madison, WI, USA), and the recombinant proteins were expressed by cold shock at 15°C for 30 min with 0.2 mM of IPTG. The harvested cells were disrupted by sonication (4°C, 50W, 5 s) using a Sonicator (Ohtake Works, Tokyo, Japan) three times, and the supernatant was adsorbed on a Talon cobalt-chelating column (Takara Bio, Shiga, Japan). To remove the fused ZZ domain and hexa histidine-tag, the column was washed using a wash buffer (50 mM Tris-HCl pH 7.0, 1 M NaCl, 15 mM imidazole), then on-column digestion reaction using PreScission protease (GE Healthcare, Piscataway, NJ, USA) was conducted overnight at 4°C for 12 h, and the protein eluted using an elution buffer (50 mM Tris-HCl pH 7.0, 150 mM NaCl, 1 mM EDTA, 1 mM DTT). Protein concentration was measured using Bio-Rad Protein Assay Dye Reagent (Bio-Rad, Hercules, CA, USA) with bovine serum albumin Fraction V (Sigma-Aldrich, St. Louis, MO, USA) as a standard. The homogeneity and concentration estimated by protein assay was confirmed by SDS-PAGE using 10% separation gel with CBB staining (Extended Data Fig. 4). We tested the stability of each recombinant protein by leaving the solution on ice, and confirmed that the luminescence activity had not decreased significantly decreased in the order of total light intensities (50-100%) for all proteins even after 12 h (data not shown). This suggests that enzymatic inactivation of the ancestral proteins during purification process was negligible. The purified protein was stored at −80°C until use.

#### Measurement of the luminescence spectra

Luminescence spectra was measured using a spectrophotometer AB-1850 (ATTO, Tokyo, Japan). Spectral sensitivity was calibrated. A 2.5 μg or 5 μg of purified recombinant luciferase was mixed with 500 μM D-luciferin, 4 mM ATP, and 8 mM MgCl_2_ in 0.1 M Tris-HCl buffer pH 8.0. Exposure time was 1-2 min. *Photinus pyralis* luciferase (PpyLuc1) was purchased from Sigma-Aldrich. Because of its very low luminescence activity, 10 μg of the protein was used for measuring the spectrum of AncElat.

#### Measurement of the luminescence intensities

Luminescence intensity was measured using a luminometer Centro LB960 (Berthold). Luminescence reaction was initiated by injecting the 50 μl of the purified recombinant luciferase (430 ng) into 50 μl of the mixture of D-luciferin (final conc. 10 μM), ATP (final conc. 100 μM) and MgCl_2_ (final conc. 5 mM) in Tris-HCl buffer pH 7.8 (final 50 mM). Light intensity was measured for 30 s after injection.

#### Measurement of the ACS activities

Thioesterification activity was determined using UV–vis spectrophotometry (Eppendorf, BioSpectrometer kinetic). In this assay, we measured the initial rate of AMP formation by coupling the thioesterification reaction with adenylate kinase, pyruvate kinase, and lactate dehydrogenase and monitoring the oxidation of NADH at 340 nm (6220 M^-1^ cm^-1^)^10^. The standard reaction mixture for this assay contained 100 mM Tris-HCl (pH 8.0), 10 mM ATP, 10 mM MgCl_2_, 0.35 mM lauric acid (C12:0) substrate, 2 mM CoASH, 1 mM phosphoenolpyruvic acid, 0.4 mM NADH, 40 μg/ml adenylate kinase, 20 μg/ml pyruvate kinase, 20 μg/ml lactate dehydrogenase, and 10 μg of purified recombinant luciferase. The total volume was brought to 500 μl. The mixture containing all components except for the luciferase was incubated at room temperature (27 ± 2°C) for 10 min. The reaction was then initiated by addition of the enzyme and the data collected at 5 s intervals for 10 min.

#### Crystal structure analyses

The structures of AncLamp were determined by X-ray crystallography. The AncLamp crystals were grown by the hanging drop vapor diffusion method, under conditions using 0.1 M trisodium citrate buffer (pH 5.5) containing 20 % (*w*/*v*) PEG3000 as a 0.5 ml reservoir, and a mixture of 2 μl of reservoir solution and 2 μl of protein solution in 50 mM Tris-HCl (pH 8.0) buffer, containing 1% (*w*/*v*) AncLamp in the hanging drop. To co-crystallize the protein with reaction-intermediate analog or substrates, 0.2 mM 5’-*O*-[*N*-(dehydroluciferyl)-sulfamoyl]adenosine (DLSA), or 0.2 mM ATP and 0.2 mM D-luciferin were added to the protein solutions. All crystals were grown at 18°C for a few weeks.

X-ray diffraction data were collected from loop-mounted crystals under cryogenic conditions, with a CCD detector Quantum315 (ADSC) at BL38B1 or Eiger4M at BL26B2 in SPring-8 (Hyogo, Japan). The crystals were soaked for 10–30 s in the corresponding crystal growth buffer, containing 15% (*v*/*v*) 2-methyl-2,4-pentanediol (MPD) for cryoprotection. The diffraction images were processed with the MOSFLM program^11, 12^.

The crystal structures were solved by the molecular replacement method, using the Phaser-MR application of PHENIX^13^. The crystal structure of *L. cruciata* LcLuc1 (PDB code 2d1s) was used for a search model. The model refinements were conducted by using COOT and the phenix.refine application of PHENIX^12, 14^. The quality of the models was evaluated with the PROCHECK program^15^. The crystallographic parameters, data collection and refinement statistics, and PDB codes are summarized in Extended Data Table 1 The atomic coordinates and structure factors of AncLamp-DLSA and AncLamp-D-luciferin-ATP complexes have been deposited in the Protein Data Bank, with the accession codes 6K4C and 6K4D, respectively. The molecular graphics were prepared with CHIMERA^16^.

### DISCUSSION

#### Crystal structure of AncLamp with DLSA or D-luciferin and ATP

The ancestral firefly luciferase AncLamp was co-crystallized with the reaction-intermediate analog 5’-*O*-[*N*-(dehydroluciferyl)-sulfamoyl]adenosine (DLSA), and the crystal structure was determined to a 2.1 Å resolution (Fig. 2 and Extended Data Table 1). The overall structure of AncLamp was well conserved with LcLuc1 (PDB 2d1s), which shows 79% amino acid sequence identity and 1.64 Å root mean square deviation for 535 C*α* atoms with AncLamp^17^. The DLSA molecule was in close contact (at least one atom of amino acid residue existed less than 3.5 Å from DLSA atoms) with His244, Phe246, Thr250, Ser313, Pro317, Gly338, Tyr339, Gly340, Leu341, Thr342, Thr361, Asp421, Ile433, Arg436, and Lys528 in the AncLamp-DLSA complex structure. These substrate-binding residues are generally conserved with LcLuc1. The conformations of DLSA molecule were nearly identical between AncLamp and LcLuc1 as showing 0.38 Å root mean square deviation for 40 DLSA atoms in the superposed structures (Extended Data Fig. 8a).

A noticeable difference in DLSA interaction was observed in proximity of the residues mutated between AncLamp and LcLuc1, namely, Ile240 (Val243 in LcLuc1) and Leu285 (Ile288). Mainly due to the packing of C*δ*1 atom of Ile240 and C*δ*2 atom of Leu285, the phenyl ring of Phe246 was slightly extruded toward the substrate-binding cavity. The distance from the C*ε*1 atom of Phe246 (Phe249 of LcLuc1) residue to the thiazole plane of DLSA (defined by atoms C8 - S9 - C4) is 3.18 Å and 3.43 Å in the AncLamp and LcLuc1 structures, respectively (Fig. 2b). Consequently, the space for luciferyl-moiety appeared to be narrower in AncLamp structure. The site-directed mutagenesis and structure analyses of LcLuc1 in the previous study revealed that elimination of a space for thermal relaxation of luciferyl moiety was one of the most significant factors for a blue-shift in the luminescence wavelength^17^. Thus, the crystal structure of AncLamp implied that the blue-shifted luminescence in the ancestral luciferase would also be explained by the spatial constraint on the luciferyl moiety to reduce energy loss through thermal relaxation.

The ancestral firefly luciferase AncLamp was also co-crystallized with genuine substrates, namely ATP and D-luciferin, and the crystal structure was determined to a Å resolution. Although the protein structure was almost identical to that of AncLamp-DLSA complex, the electron density map at the active site demonstrated an intriguing feature, suggesting the presence of the reaction intermediate, most probably a luciferyl-AMP. The conformation of luciferyl-AMP was very close to that of DLSA as showing 0.42 Å root mean square deviation for 40 atoms (Extended Data Fig. 8b). The conformations of the substrate-interacting residues are also similar between the complex structures DLSA and luciferyl-AMP complex (Extended Data Fig. 8a). Thus, the space -elimination for luciferyl moiety, which was observed for the intermediate analog (DLSA) complex, appeared to be also consistent for genuine intermediate molecule.

Furthermore, significant residual density was observed after building the luciferyl-AMP model, and it was interpreted to be unreacted D-luciferin. When a *Fo* – *Fc* electron density map was synthesized by omitting the luciferyl-AMP molecule, few significant peaks of residual densities, which did not correspond to luciferyl-AMP atoms, were observed. These residual densities fit well to sulfur or oxygen atoms when unreacted D-luciferin was built into the substrate-binding cavity (Extended Data Fig. 8c). Unexpectedly, the unreacted D-luciferin was trapped in the substrate-binding cavity upside-down for reaction, apparently showing unproductive configuration. According to the PDB, no structure of luciferase complexed with D-luciferin or with D-luciferin and ATP has been reported so far. Therefore, significance of the observed D-luciferin configuration for luciferase is not conclusive at this point of time. However, it could be speculated that early luciferases were not highly adapted to D-luciferin as a specific substrate as luciferases have been evolved from fatty acyl-CoA synthetases, which accept various fatty acid substrates. As demonstrated by the acyl-CoA synthetase activity assay, which is detailed below, firefly luciferases have evolved a specificity for D-luciferin. The observed unproductive configuration might be a result of immature specificity of AncLamp toward D-luciferin.

#### ACS activity

Our previous reports have suggested that firefly luciferases and the orthologues in beetles and *Drosophila* possess acyl-CoA synthetase (ACS) activity, and the most preferable substrate was lauric acid^18^. Based on these results, we have hypothesized that beetle luciferase originated from a fatty acyl-CoA synthetase (presumably medium-chain fatty acyl-CoA synthetase)^18, 19^. In this context, we measured the ACS activities of the purified ancestral luciferases using lauric acid as substrate. The results showed that ACS activities of the ancestral luciferases were comparable to these of extant luciferases (*L. cruciata* LcLuc1, LcLuc2 and *P. pyralis* PpyLuc1), except for the case in AncElat which exhibited about 10 times higher in activity (Extended Data Fig. 7). This conclusion is in strikingly agreement with our ‘ACS origin’ hypothesis of beetle luciferase. Furthermore, it also fits to the evolutionary ‘trade-off’ model of new enzyme, in which original catalytic function of an enzyme is decreased during the process of acquiring new catalytic function^20^. Indeed, we previously suggested that the ‘trade-off’ model fits the evolution of beetle luciferase by site-directed mutagenesis studies of beetle fatty acyl-CoA synthetase^21^. In the aforementioned paper, we constructed all combinations of the mutants for 3 amino acids near active site in AbLL, an ACS of non-luminous click beetle, and measured both luciferase and ACS activities. The results have showed negative trade-off of these activities among mutants, that is, mutants having higher ACS activity exhibited lower luciferase activity, and *vice versa*^21^. The point is that firefly luciferase and ACS are both ‘promiscuous’ for substrate specificity and reaction property; native firefly luciferases have ACS activity to various fatty acids^18, 22^ and some native ACSs have weak but significant luciferase activity to D-luciferin^21^, that warrants evolvability of new enzyme^20^. Recently, it has been reported that CG6178, an ACS in *Drosophila* and AbLL exhibited over 1,000 times higher luminescence activity with synthetic luciferin analogues, CycLuc2 and CycLuc12, compared to those with native D-luciferin^23, 24^. These results support that ACSs possess ‘latent’ luciferase activity with the potential to evolve further. Our present study showed that AncElat, the protein of the common ancestor between cantharoids and elaterids which was probably non-bioluminescent^19^, had high ACS activity with low luciferase activity. On the other hand, AncCanth, the protein of the common ancestor of cantharoids which was probably bioluminescent^19^, emitted light more than 1,000 times greater than AncElat (Extended Data Fig. 5b) and had ACS activity about 10 times lower than AncElat (Extended Data Fig. 7), which demonstrated a negative trade-off process for neofunctionalization. Firefly genome analysis showed that firefly luciferase evolved by tandem gene duplications of ACS and subsequent acquisition of the luciferase activity in a redundant gene^19^. Probably the gene duplication of ACS occurred in ancestral non-bioluminescent beetle leading to the evolution of a new function, luminescence activity, without suffering any trade-off disadvantages.

#### Evolutionary scenario of beetle luciferase

Based on the biochemical analyses of the recombinant proteins of the ancestral firefly luciferases, we suggest that the last common ancestor firefly (AncLamp) emitted green light as a warning display of toxicity/distastefulness. Based upon this supposition, we propose an evolutionary stepwise scenario for firefly luciferase using the enzymatic properties of seven ancestral luciferases ranging from the pre-bioluminescent dark period to the divergent success of the family Lampyridae.

(i) In the beginning, prior to the divergence of the ‘cantharoid’ group and elaterid lineages, the luminescence activity of the ‘proto-luciferase’ (AncElat) was rudimentary, thus the faint reddish luminescence may have been non-functional, biologically. This supports the hypothesis of the parallel emergence of bioluminescence in the ‘cantharoid’ group and Elateridae^19^. We believe the reddish trace luminescence activity in AncElat represents a prototypical state unable to produce light with high intensity and a shorter wavelength (higher energy), as previously suggested^25, 26^. In contrast, this proto-luciferase was a highly specialized fatty acyl-CoA synthetase, its original function.

(ii) When the luciferase acquired substantial luminescence ability in the ‘cantharoid’ lineage, AncCanth elicited yellow, almost orange light, probably due to the evolution of an exposed active site. The biological function of the ancient beetle bioluminescence about 100 Mya was unknown, but it may have functioned as a supplementary aposematic warning for toxicity/distastefulness at night^27^.

(iii) The common ancestor of extant fireflies was probably nocturnal, and the luminescence produced by the luciferase AncLamp developed into a primary biological aposematic display function, accompanied by an increase in brightness and shifting in colour to green. This change was enabled by D-luciferin being tightly bound, as demonstrated by the crystal structures. The ancestral firefly shifted in behaviour from diurnal to nocturnal.

(iv) Gene expression profiles suggest that Luc1-type luciferase is responsible for the luminescence of the lantern in larvae, pupae and adults, while Luc2-type luciferase is responsible for the dim glow of oviposited eggs, the pupal body, and the ovaries^28, 29^. After gene duplication, both Luc1 and Luc2 initially emitted in the green spectrum (AncLuc1 and AncLuc2) as seen in the extant *Pyrocoelia atripennis*^29^.

(v) In both lineages of Lampyrinae and Luciolinae, some species independently began to adapt to a twilight environment for mating behaviour. Now the crepuscular fireflies needed to differentiate mate luminescence from the background noise of green foliage wherein lantern emissions became more red-shifted toward yellow (AncLampn1 and AncLucin1) as seen in extant *Photinus pyralis* and *Luciola parvula*^30^. Interestingly, the spectral sensitivities of crepuscular fireflies match their own yellowish light spectra to receive the conspecific luminescence effectively^31, 32^.

(vi) In contrast to the colourful evolution of Luc1-type luciferase, extant Luc2-type luciferase took over the original green luminescence (AncLuc2) for warning displays in egg and pupal stages in both diurnal and nocturnal fireflies.

## Extended Data

**Extended Data Table 1.** X-ray data collection and refinement statistics^a^.

| Protein<sup>b</sup> | AncLamp - DLSA<br>complex<br>(6K4C) | AncLamp – ATP –<br>D-luciferin complex<br>(6K4D) |
| --- | --- | --- |
| Space group | *C222<sub>1</sub>* | *C222<sub>1</sub>* |
| Cell constants (a, b, c Å) | 48.3, 123.4, 177.4 | 48.8, 123.7, 177.0 |
| Wavelength (Å) | 1.0000 | 1.0000 |
| Resolution range - data<br>collection (Å) | 20.00 - 2.10 (2.21 - 2.10) | 20.00 - 1.70 (1.79 - 1.70) |
| R<sub>sym</sub> | 0.032 (0.073) | 0.014 (0.030) |
| Completeness (%) | 99.7 (100.0) | 96.3 (95.4) |
| $\langle I/\sigma I \rangle$ | 14.0 (6.9) | 75.6 (22.5) |
| Resolution range -<br>refinement (Å) | 20.00 - 2.10 (2.17 - 2.10) | 20.00 - 1.70 (1.73 - 1.70) |
| No. reflections | 31197(3317) | 55829 (2520) |
| R<sub>cryst</sub> | 0.208 (0.266) | 0.163 (0.226) |
| R<sub>free</sub> | 0.244 (0.328) | 0.189 (0.280) |
| RMSD length (Å) | 0.007 | 0.017 |
| RMSD angle (°) | 0.98 | 1.22 |

<sup>a</sup> Values for the highest resolution bin are in parentheses.<sup>b</sup> In parentheses PDB accession codes.

**Extended Data Table 2.**
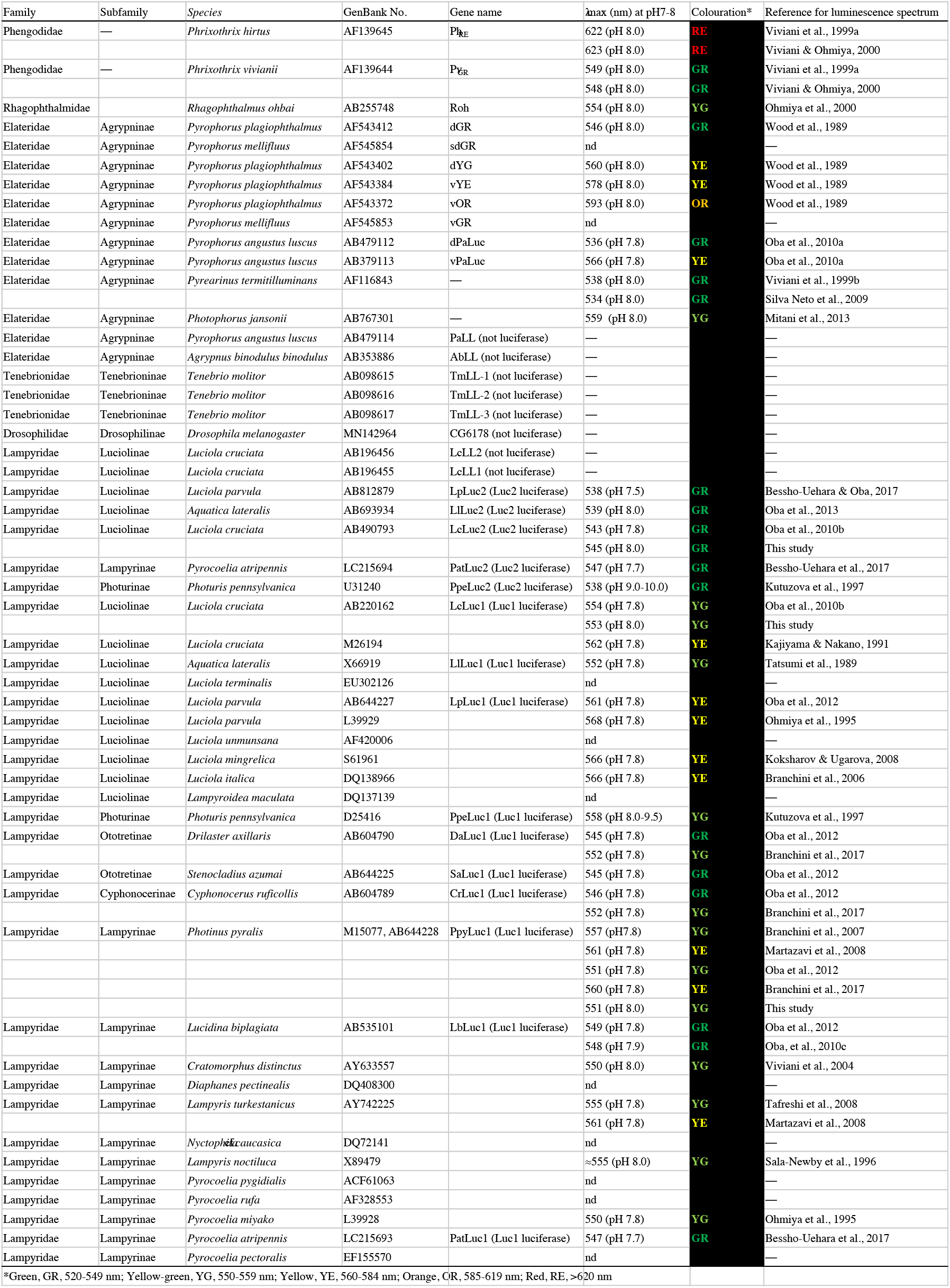
Extant luciferase genes and the luminescence spectral maximum.

**Extended Data Fig. 1.**
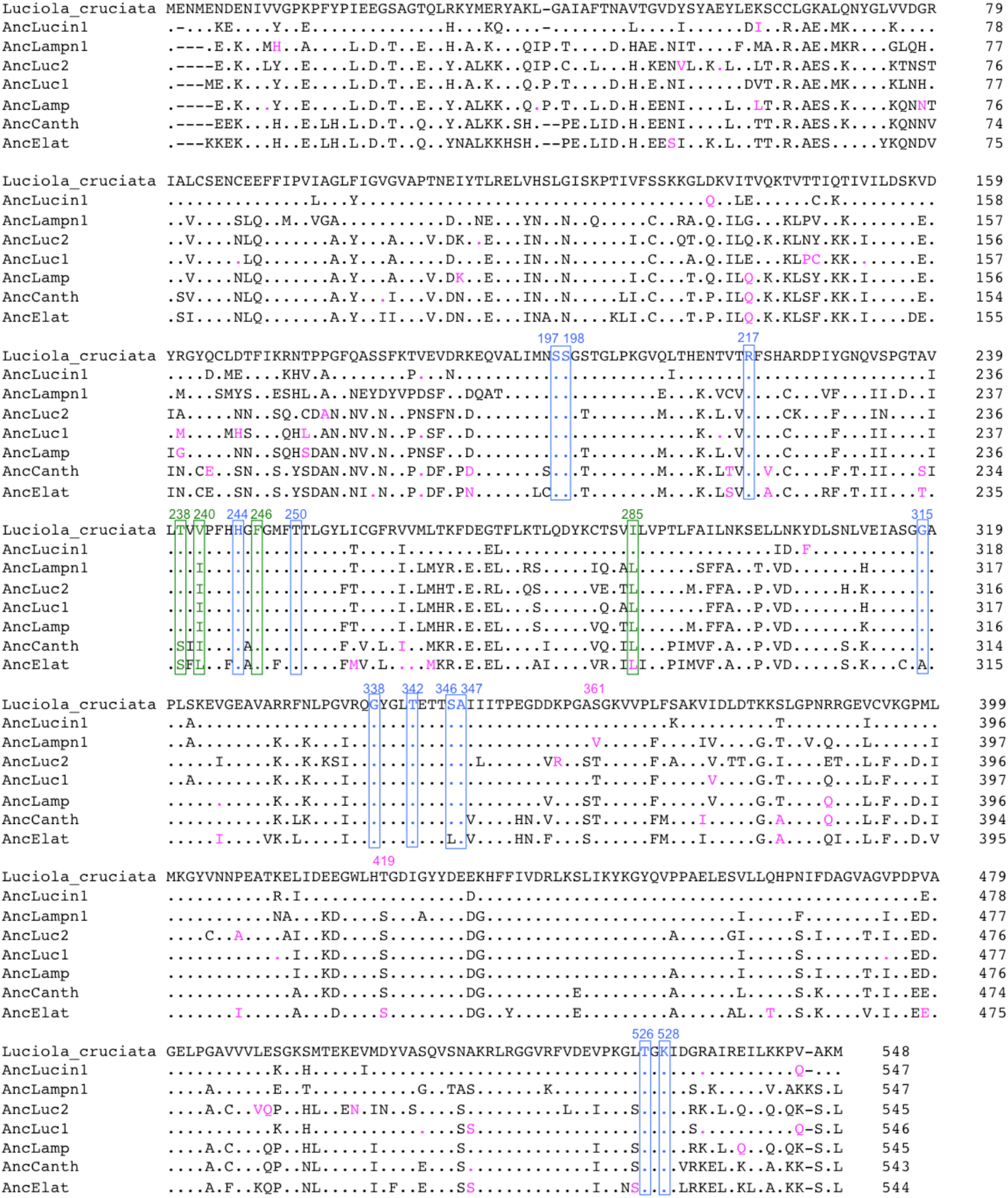
Multiple-sequence alignment of ancestral luciferases and *Luciola cruciata* LcLuc1. The amino acid residues of the ancestral luciferases, which are identical to LcLuc1, are shown as a dot. The active site residues (Ser197, Ser198, Arg217, His244, Thr250, Gly315, Gly338, Thr342, Ser346, Ala347, Thr526, and Lys528 reference the positions in AncLamp) (Nakatsu et al., 2006) are outlined in blue. The predicted sites relevant to the AncLamp luminescence spectra (Thr238, Ile240, Phe246, and Leu285) are outlined in green (shown on structure in Fig. 2). The more ambiguously identified sites (posterior probability less than 0.1 higher than the second most probable amino acid) are shown in magenta (depicted on structures in Extended Data Fig. 3).

**Extended Data Fig. 2.**
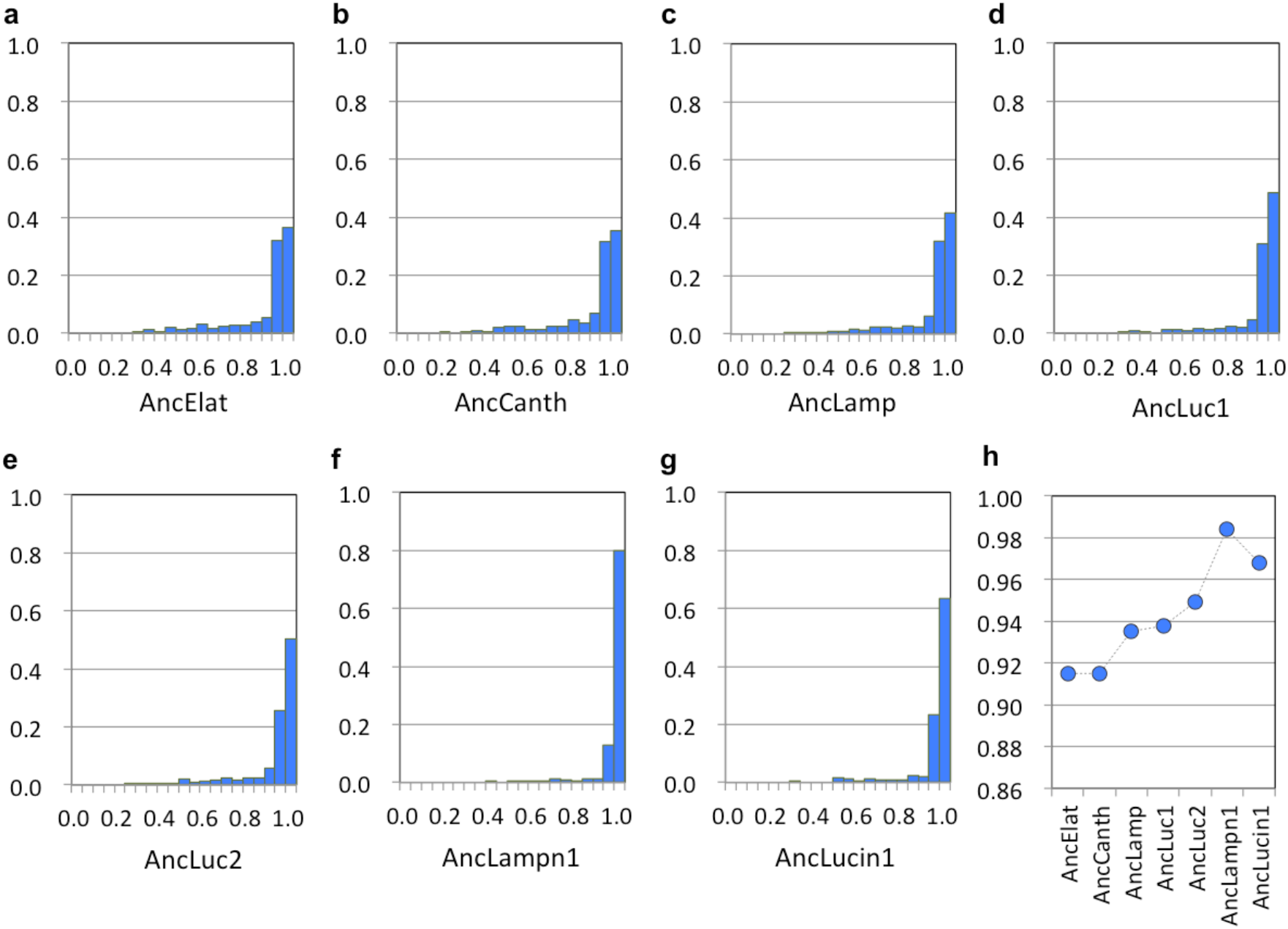
Posterior probability distributions of the residue sites of **a**, AncElat, **b**, AncCanth, **c**, AncLamp, **d**, AncLuc1, **e**, AncLuc2, **f**, AncLampn1, and **g**, AncLucin1 (G). Horizontal and vertical axes show posterior probability bins and fractional frequency of inferred sites, respectively. **h**, Average posterior probabilities are plotted over the ancestral luciferases.

**Extended Data Fig. 3.**
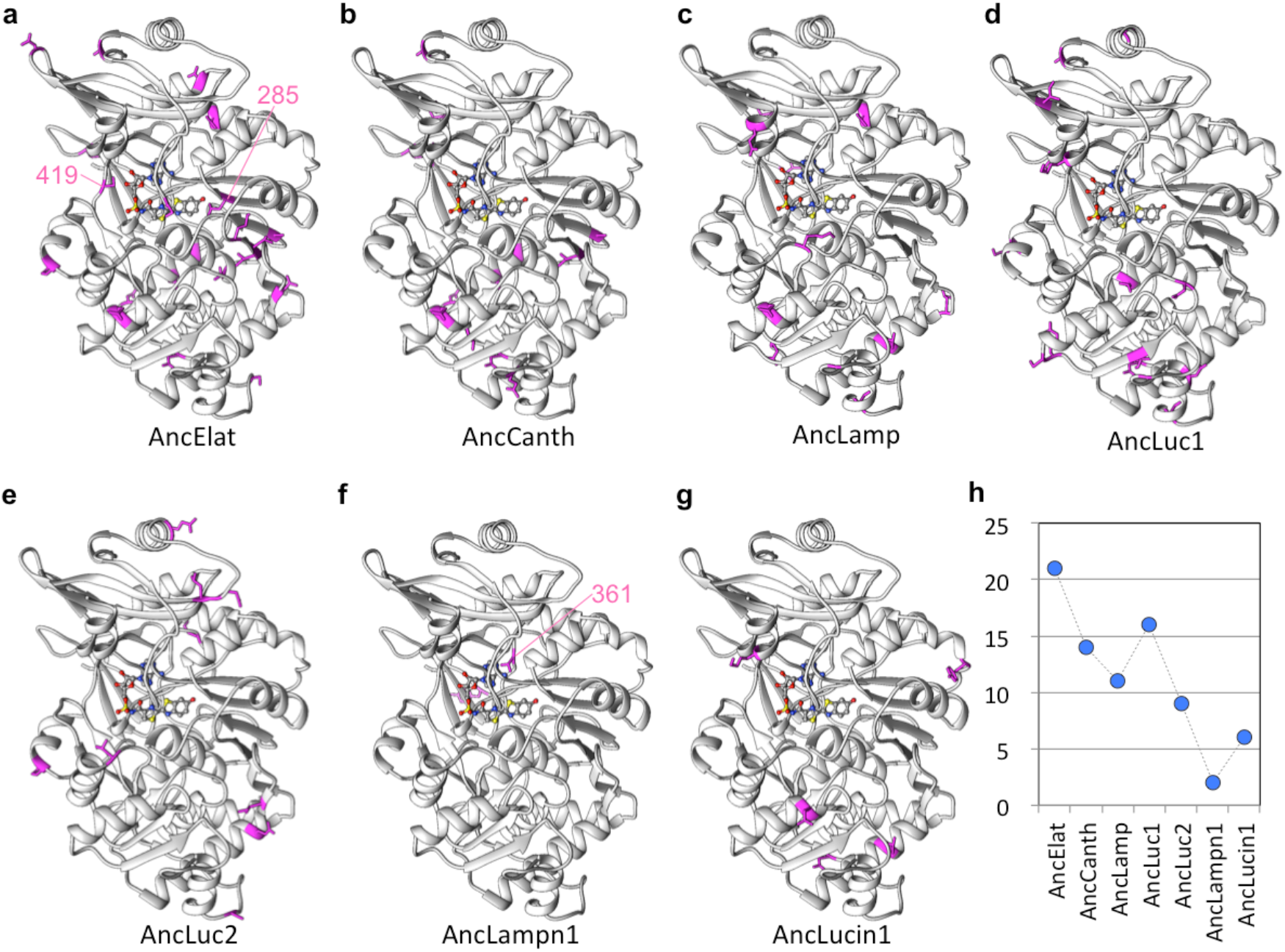
Ambiguous residues, for which the posterior probability is less than 0.1 higher than the second most probable amino acid, are shown in magenta on **a**, AncElat, **b**, AncCanth, **c**, AncLamp, **d**, AncLuc1, **e**, AncLuc2, **f**, AncLampn1, and **g**, AncLucin1 structure models. The sites that might have direct interaction with the active site residues, namely Val361 in AncLampn1, and Leu285 & Ser419 in AncElat (also indicated in Extended Data Fig. 1) are denoted with residue number. **h**, Total numbers of ambiguous residues are plotted over the ancestral luciferases.

**Extended Data Fig. 4.**
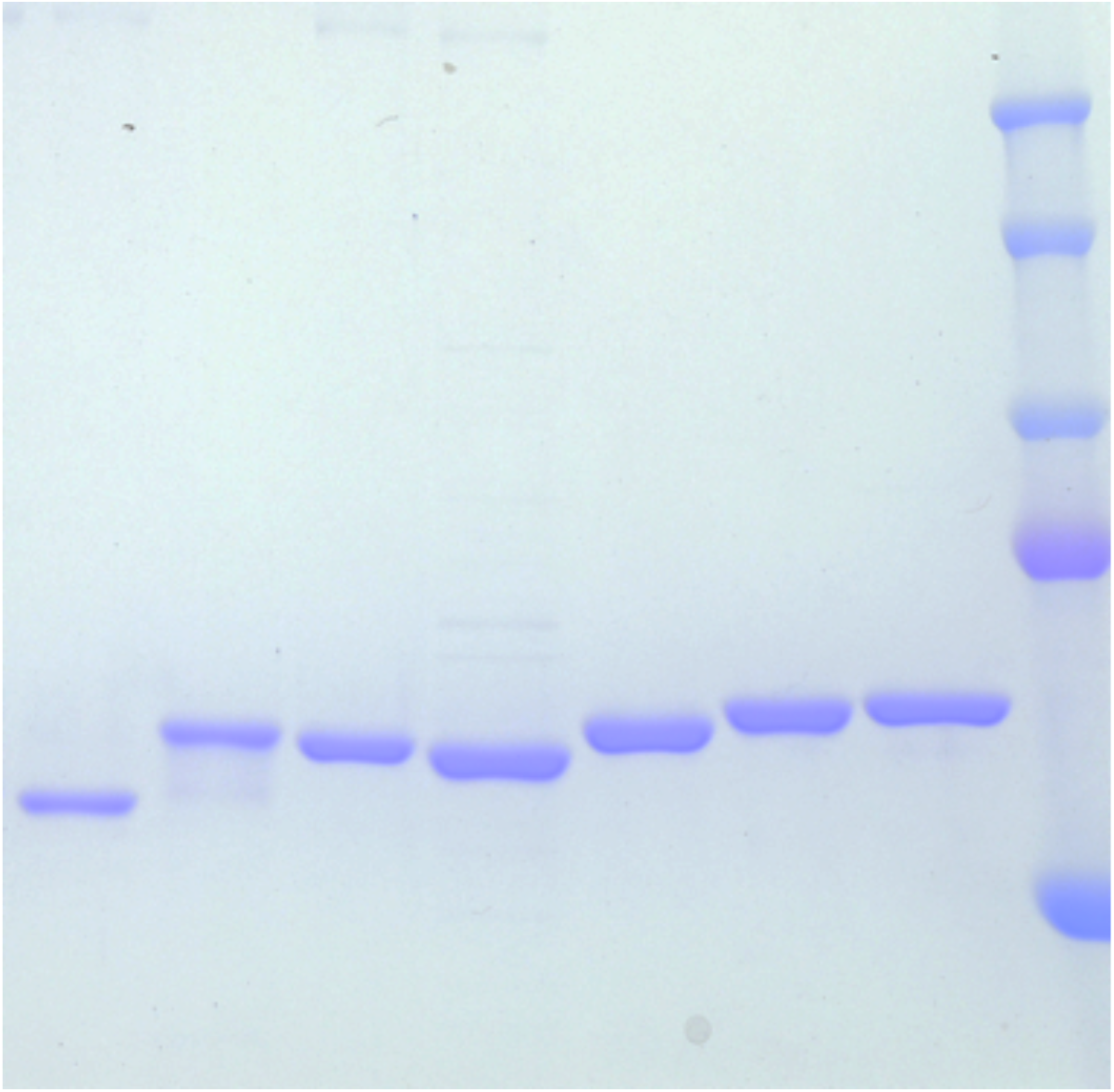
SDS-PAGE gel stained by CBB. 430 ng of each protein was loaded. From left: AncElat, AncCanth, AncLamp, AncLuc2, AncLuc1, AncLucin1, AncLampn1, and molecular marker (from top, 250, 150, 100, 75, 50 kDa; Precision Plus Protein Dual Standards, Bio-Rad).

**Extended Data Fig. 5.**
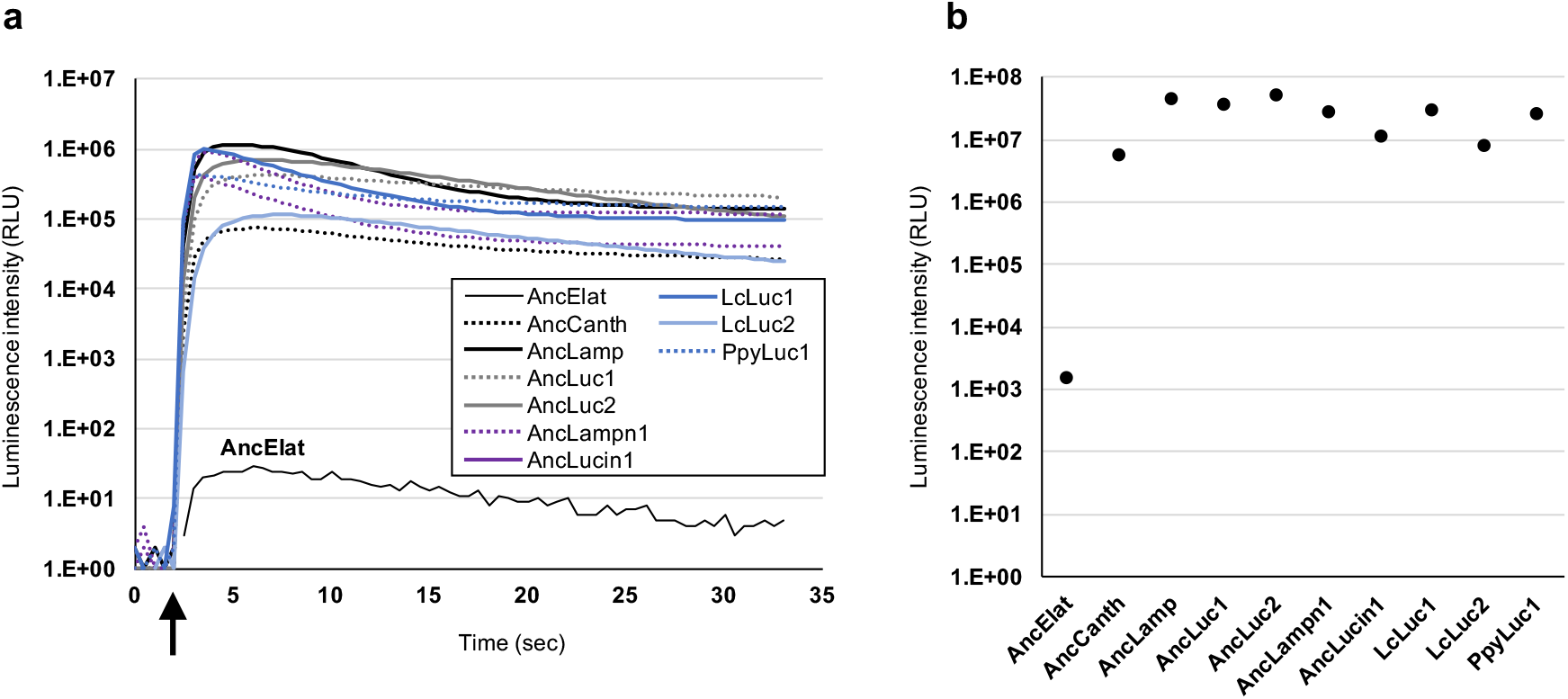
**a**, Time-course luminescence of 7 recombinant ancestral and 3 extant luciferases. Luciferin, ATP and other components were injected into luciferase solution after 2 s (arrow). **b**, Integration from 2 s to 32 s. The values are shown by relative light unit (RLU).

**Extended Data Fig. 6.**
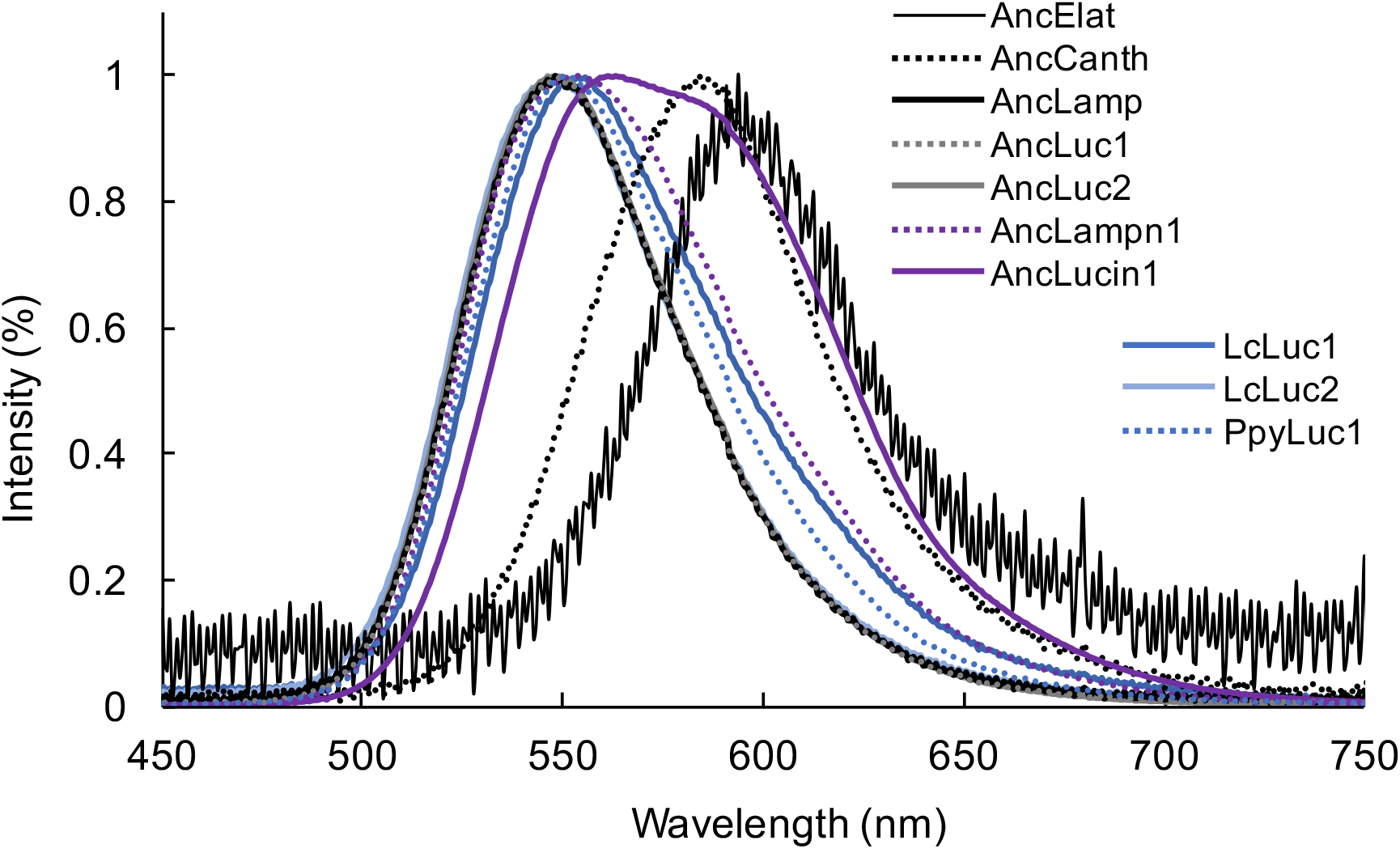
Luminescent spectra of 7 recombinant ancestral luciferases and 3 extant firefly luciferases.

**Extended Data Fig. 7.**
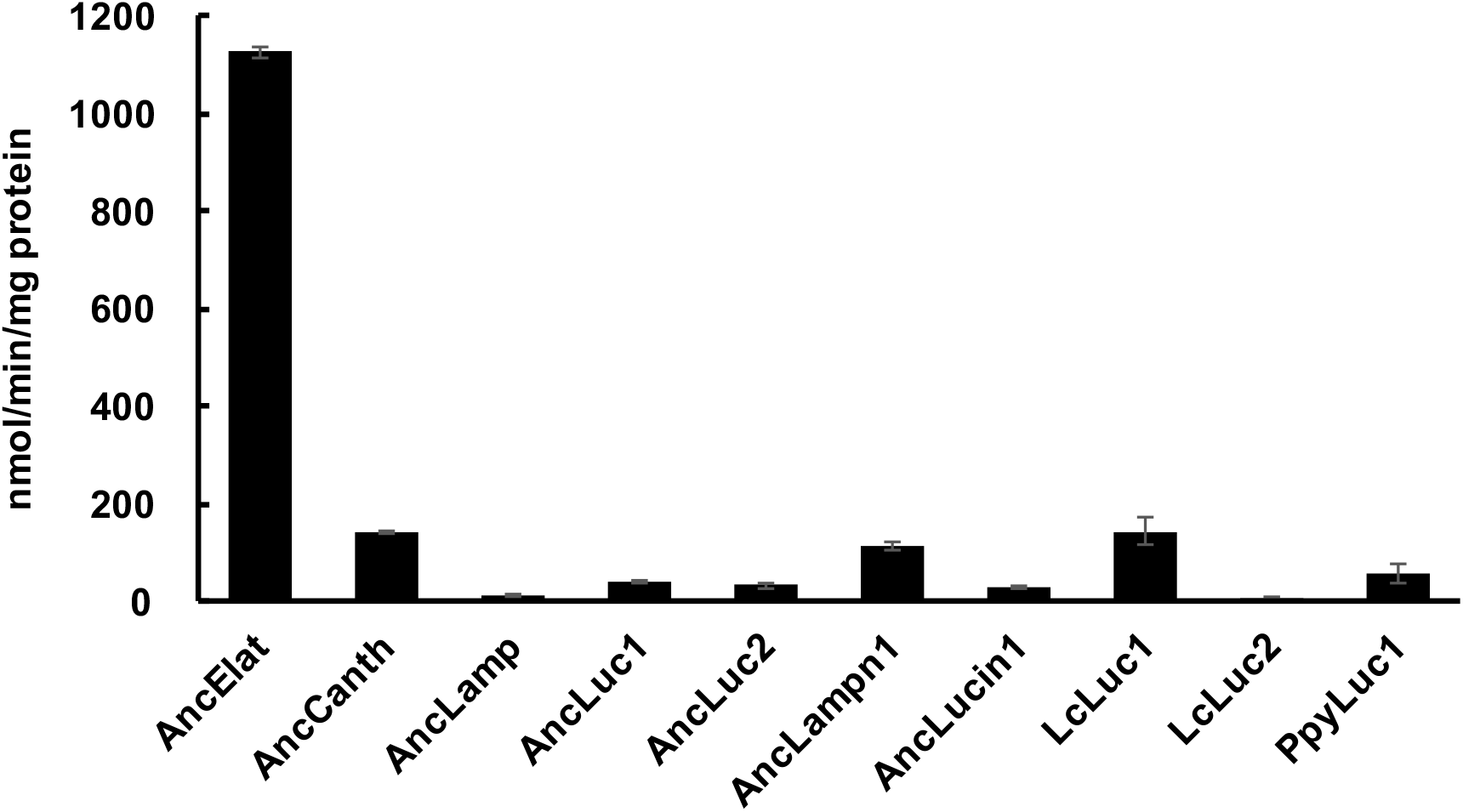
Acyl-CoA synthetic activities of 7 ancestral luciferases and 3 extant firefly luciferases to lauric acid are shown. Bars represent SE of the means (*n* = 3-6).

**Extended Data Fig. 8.**
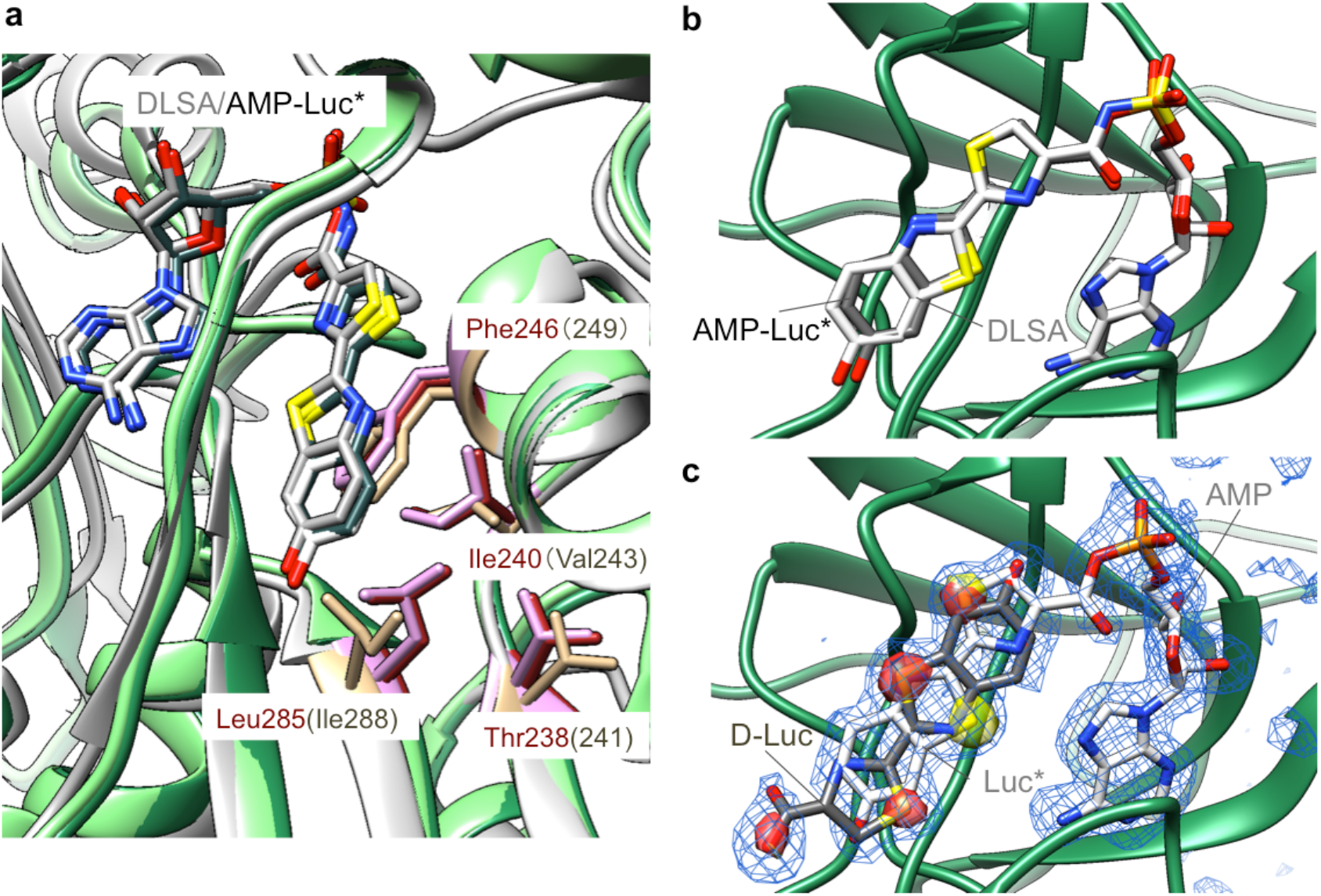
**a**, Detail of the substrate-binding site of superposed AncLamp - DLSA (main chain and side chain are colored green and red, respectively), LcLuc1 - DLSA (white and light brown), and AncLamp - substrates (D-luciferin/ATP) (light green and pink) complex structures. The residues differing between AncLamp and LcLuc1, Ile240 (Val243 in LcLuc1) and Leu285 (Ile288), and those in close contact with these residues, Phe246 (Phe249) and Thr238 (Thr241) are shown in stick models. The molecules of the DLSA and probable intermediate, luciferyl-AMP (Luc*-AMP), are shown as stick models colored by element (carbon atoms are coloured in white, grey, and slate grey for AncLamp – DLSA, LcLuc1 – DLSA, and AncLamp - substrates complex structures, respectively). **b**, The structures of AncLamp-DLSA complex and the AncLamp – substrates (D-luciferin/ATP) complex are superposed. The carbon atoms of DLSA and probable reaction intermediate luciferyl-AMP (Luc*-AMP) are shown in white and grey, respectively. **c**, The all ligand omitting *Fo* – *Fc* map is shown in blue mesh (contoured at 10 *σ*). Also the *Fo* – *Fc* D-Luc omitting map (red, 30 *σ*), and the Luc* omitting map (yellow, 30 *σ*) are superposed. The probable non-reactive D-luciferin (D-Luc, carbon atoms are shown in gray) and AMP and luciferyl moiety in the probable reaction intermediate (Luc*, carbon atoms are shown in white) are indicted with stick models.

**Extended Data Fig. 9.**
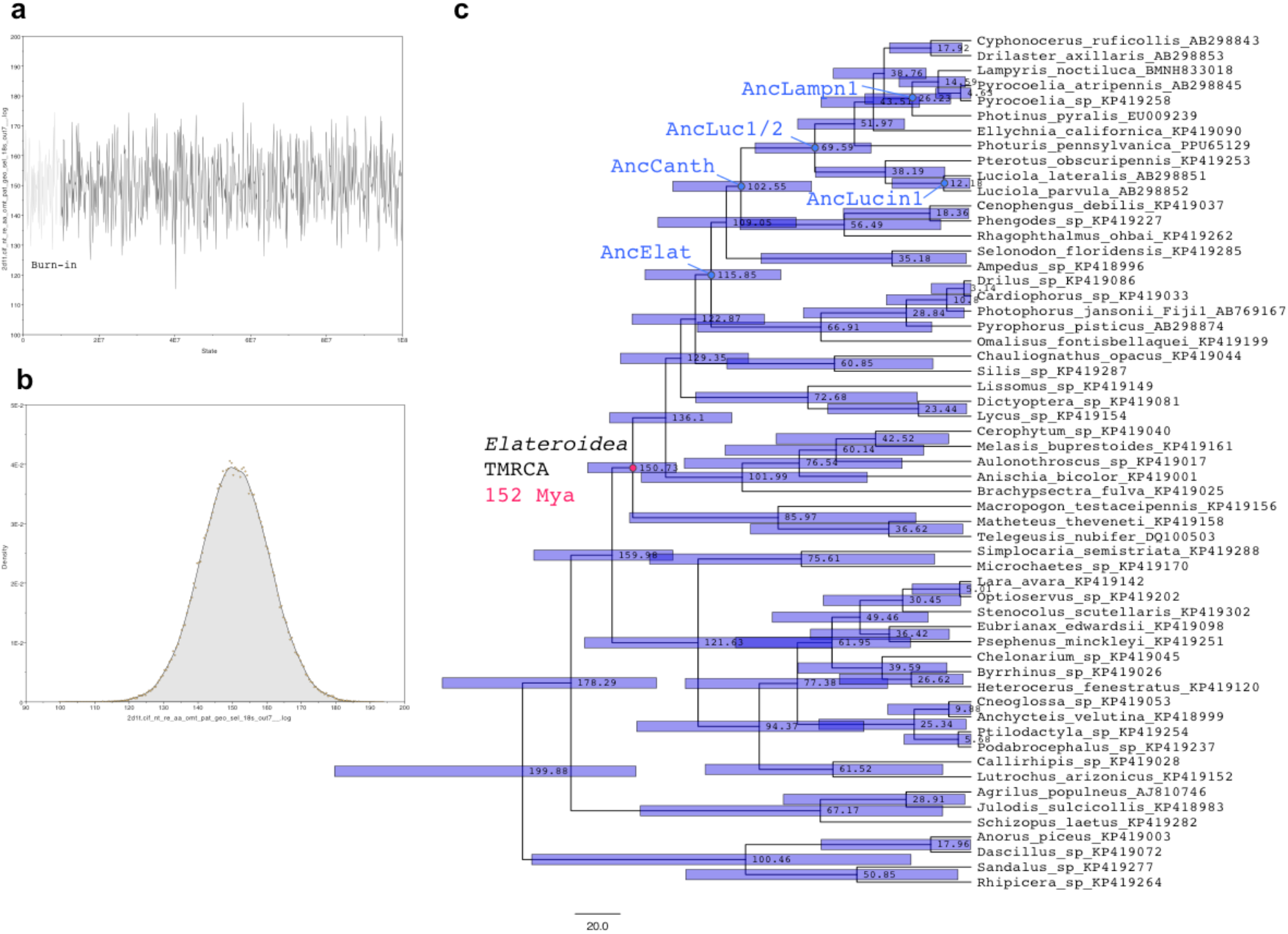
**a**, The trajectory and **b**, Distribution of Elateroidea common ancestor geological age during MCMC simulation for 10^8^ iterations. The initial 10^7^ iterations are burn-in process. **c**, The phylogeny of Elateriformia species. The leaf nodes are indicated with species names and GenBank accession number of corresponding 18S rRNA gene sequence. The median of estimated geological age and 95% HPD (purple bar) are indicted on each node. The reference Elateroidea TMRCA (McKenna et al., 2015) is highlighted with a magenta circle, and the nodes corresponding to ancestral luciferases are shown with blue circles.

